# PathoEye: a deep learning framework for the whole-slide image analysis of skin tissue

**DOI:** 10.1101/2025.06.16.659818

**Authors:** Yusen Lin, Feiyan Lin, Yongjun Zhang, Jiayu Wen, Guomin Li, Xinquan Zeng, Hang Sun, Hang Jiang, Teng Yan, Ruzheng Xue, Hao Sun, Bin Yang, Jiajian Zhou

## Abstract

**Objective:** To provide an interpretable computational framework for the examination of whole-slide images (WSI) in skin biopsies, PathoEye focuses on the dermis-epidermis junctional (DEJ) areas, also known as the basement membrane zone (BMZ), to enrich the pathological features of various skin diseases.

**Methods:** We presented PathoEye for WSI analysis in dermatology, which integrates epidermis-guided sampling, deep learning and radiomics. It enables unsupervised semantic segmentation of the BMZ and extracts distinct features associated with various skin conditions.

**Results:** PathoEye performs comparably with existing methods in binary classification tasks, while outperforming them in multi-classification tasks involving different skin conditions. It enables the investigation of histopathological aberrations in aged skin compared with young skin. Additionally, it highlighted the texture changes in the BMZ of young skin compared with aged skin. Further experimental analyses revealed that senescence cells were enriched in the BMZ, and the turnover of basement membrane (BM) components, including COL17A1, COL4A2, and ITGA6, was increased in aged skin.

**Conclusion:** PathoEye is a comprehensive tool for characterizing the unique features of different skin conditions associated with BMZ, thereby ensuring its potential applications in skin disease diagnosis and treatment planning.

## Introduction

Histological examination of skin biopsy slides is the gold standard for diagnosing most skin diseases in clinical settings ^**1,2**^. Advances in technology have enabled the accumulation and use of high-resolution whole-slide images (WSIs) in routine clinical practice ^**2–5**^. Currently, whole-slide imaging systems allow pathologists and researchers to visually inspect only a limited number of WSIs ^**6–8**^. Nonetheless, summarizing features for comparative analysis across a large set of WSIs remains challenging: 1) precisely extracting the targeted areas of WSIs; 2) exploring novel features of histological images in an unsupervised manner ^**2,9–11**^.

Recent studies have applied various algorithms and techniques to identify and quantify different features and patterns in the WSI of skin tissues ^**12–16**^. For example, Chen et al. developed a generalized and efficient model for the hematoxylin and eosin (H&E) stained WSI analysis, enabling a fast and scalable search of a large dataset by analyzing the down-sampling patches from the region of interested (ROI) ^**17,18**^. Similarly, Zheng et al. demonstrated that FastMDP-RL can automatically detect melanoma with WSIs using segmentation of ROI ^**17**^. However, those methods only focus on the informative features derived from ROI while losing information on other compartments. Interestingly, some studies have indicated that the epidermis and the basement membrane zone (BMZ) are suitable for WSI analysis, as many cytological features are observed in these areas ^**18,19**^. Notably, the basement membrane (BM) provides tissue integrity, elasticity, and mechanical signaling among the cell niche, and aberrations in BMZ of various skin diseases are observed ^**20,21**^. Therefore, we propose that patches sampled along the BMZ in WSIs could provide valuable insights for comparative analyses of different skin conditions.

Manual annotation of the WSI of skin tissue is labor-intensive and time-consuming. Most existing methods require substantial manually curated WSIs, which limits their applications in clinical diagnosis ^**17,18,22**^. Interestingly, radiomics is the extracted quantitative metrics of medical images, which are easy to retrieve in high speech and have fewer computational resource requirements compared with traditional methods ^**23–26**^. On the other hand, the deep convolutional neural network (DCNN) and foundation model have been employed in WSI analysis by dividing the ROIs into several patches. However, these approaches struggle to summarize the features of a disease group, often resulting in less interpretable outcomes ^**13,27–29**^. Grad-CAM has been developed to visualize the informative features from DCNN models through gradient-based localization, enhancing the deep neural network more interpretable ^**30**^. Therefore, a computational framework incorporating radiomic metrics and the interpretable Grad-CAM model may facilitate the exploration of novel features in large-scale WSI studies.

In this study, we introduced a machine learning framework for WSI analysis in dermatology, called PathoEye. This framework enables unsupervised semantic segmentation of the BMZ and extracts distinct features associated with various skin conditions. It performs comparably to existing methods for binary classification of healthy versus diseased skin while demonstrating superior performance in multi-classification tasks. Notably, PathoEye quantitatively measures epidermal thickness and observes that the variance in rete ridge length decreases with aging skin. Furthermore, it identifies and emphasizes texture changes in the dermis-epidermis junction (DEJ) areas of the aged group compared to the young group. These features highlight PathoEye’s potential applications in diagnosing skin diseases and draw attention to the functional role of BMZ integrity in skin aging.

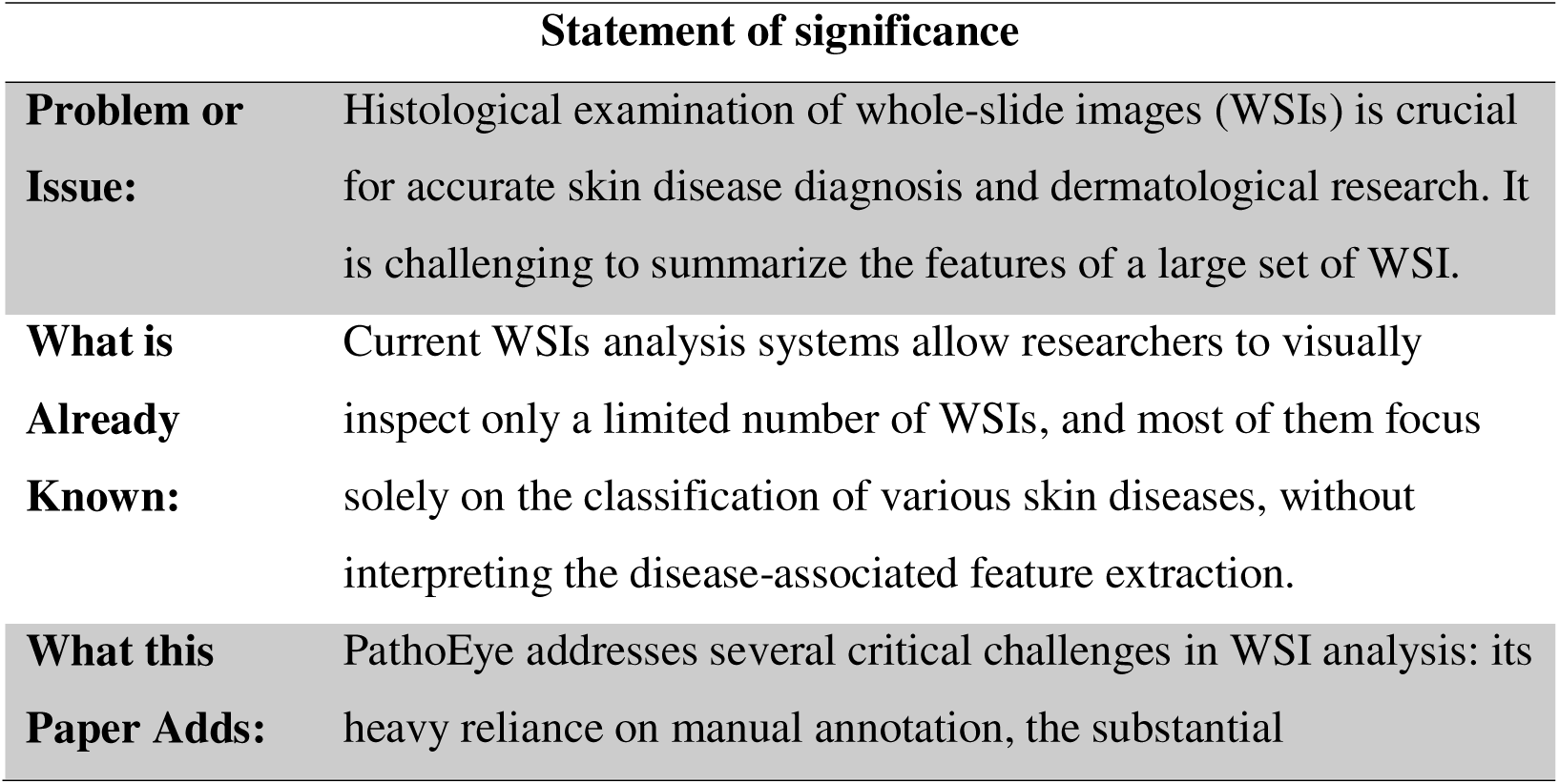
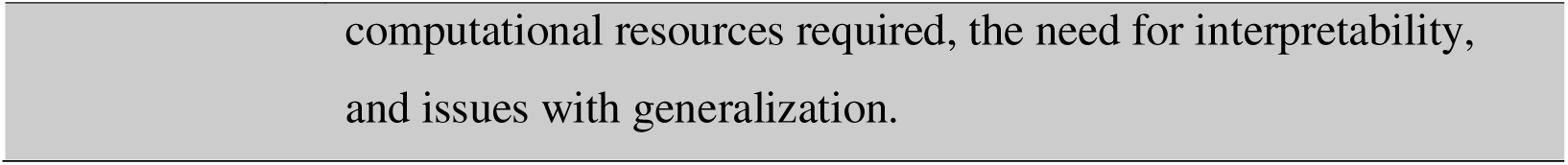

## Methods and Materials

### Data collection and sample acquisition

The WSI dataset comprises 511 exposed skin tissues and 406 non-sun-exposed skin tissues downloaded from the GTEx project ^**31**^. We defined 20-39 as the young group (n=83) and 60-79 as the aged group (n=83). Skin biopsies for Immunofluorescence (IF) and Immunohistochemistry (IHC) staining (Table S1) were obtained with ethical approval at the Dermatology Hospital of Southern Medical University (#C0225020). The H&E WSIs of 4 skin diseases are retrieved from Dermatology Hospital of Southern Medical University. All participants have signed the informed consent.

### Performance evaluation

The level 2 patches sampled from both the young and aged groups were used to evaluate the performance of PathoEye. CLAM, transMIL, and DSMIL were downloaded from GitHub and implemented under the same computational environment. We then conducted 5-fold cross-validation analyses for all four models across three classification tasks, including distinguishing aged skin tissues from the young group, differentiating diseased skin tissues from healthy ones, and classifying skin tissues under five conditions (Figures 2B and 2H). To evaluate the performance of the four methods, we assessed accuracy, receiver operating characteristic (ROC) curves, F1 scores, and the area under the precision-recall (PR) curve (AUC).

### Thickness and rete ridge score calculation

We defined EB and ES as the epidermis-dermis and epidermis-corneum boundary; The points along EB to ES or from ES to EB was used to quantitatively measure the rete ridge length and thickness, respectively (Figure 3A and Figure S3). Then, the epidermis thickness as the mean of the shortest distance of a point from ESi to EB 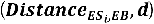 and we calculated the epidermis rete ridge score as the variance of the shortest distance of a point from EBj to ES 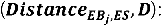

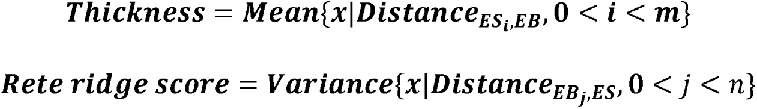

### Radiomic analysis

106 radiomic features were automatically extracted using the Pyradiomics package, utilizing level 2 or 3 patches as input ^**32**^; The features were categorized into two groups: 1) first-order statistics, and 2) higher-order texture characteristics. First-order statistics comprise 18 basic metrics that reflect the symmetry, uniformity, and local intensity distribution changes of the measured voxels. In contrast, the higher-order texture features assess the heterogeneity differences within the epidermis areas in a texture format. Then, we established a radiomic-based classifier using a random forest algorithm to classify the young and aged skin images with radiomic features. The informative features representing the differences between 2 groups were obtained.

### DCNN-based hotspot tracing module

A DCNN model is established for discriminating between young and aged skin using a pre-trained ResNet50 model as the backbone ^**33**^. Then, we applied Grad-CAM ^**30**^ to trace the defects in the aged group using the trained model. The spots in the image associated with the aged skin were extracted using the derivative of the loss value and the activation value propagated from the trained model, resulting in a heatmap representing their activation values and the derivative of the loss value in the first convolutional layer (Figure S2). The red spot represents the importance of spatial features in terms of skin aging, while the blue spot represents the less important features.

### IF and IHC staining

For IF staining, the sections were prepared and incubated with rabbit anti-p16^INK4a^ antibody (1:100 dilution; Abcam, ab108349) and the Goat Anti-Rabbit secondary antibodies (1:500 dilution; Abcam, ab150078) as described previously ^**34**^. Then, the fluorescence images were obtained with the confocal (Nikon A1+, Japan). For IHC staining, skin biopsies were fixed, sectioned and incubated with the antibodies described previously ^**35**^. Then, the WSIs were obtained using NanoZoomer (Hamamatsu, Japan). Finally, ROI intensity was analyzed using ImageJ (version 1.54f).

### Statistical analysis

Data was analyzed using Python (version 3.7) and represented as meanLJ±LJstandard deviation (S.D.). All tests were two-sided, and PLJ<LJ0.05 was considered statistically significant.

## Results

### PathoEye: a deep learning framework for WSI analysis of skin tissue

The comparative analysis of the large repositories of gigapixel WSIs requires intensive manual annotations and enormous computational resources, limiting their applications in dermatological research ^**36**^. Here, we developed PathoEye to automatically analyze the anatomy and pathological features of WSIs with less computational resources. PathoEye composites with four modules (Figure 1 and Supplementary notes): 1) epidermis extraction module; it incorporates an unsupervised segmentation algorithm InfoSeg ^**37**^ to roughly retrieve the epidermic area and apply a customized method to remove hair follicle, gland and vascular structure, resulting in level 1 images (Figure 1A and Figure S1A). 2) epidermis guided patch sampling module; it generates 512 x 512-pixel patches (term as level 2 images) along the mask representing the epidermis region in WSIs, which composite with the dermis, BM, epidermis, and stratum corneum; in the meantime, 128 x 128-pixel patches (term as level 3 images) were extracted from the level 2 images along the mask representing epidermis; they represent the details information of BMZ (Figure 1B and Figure S1B). 3) Deep Convolutional Neural Networks (DCNN) classification module; the level 2 and level 3 images are subjected to a DCNN-based model for training an unsupervised model for category classification (Figure 1C and Figure S2). 4) exploration and discovery utilities; it extracts the radiomic features of the level 2 and level 3 images using Pyradiomics ^**32**^, resulting in 106 features per image, then establishes an optimized machine learning model for classification (Figure 1D, left panel and Figure S4); on the other hand, we developed a hotspot tracing module to trace the important hotspot area of the trained model using Grad-CAM^**30**^ (Figure 1D, right panel and Figure S2), which represents the informative features for discriminating abnormal and normal skin tissues. PathoEye has three unique features: 1) it segments the skin structure of WSIs by incorporating epidermis extraction and epidermis-guided patch sampling in an unsupervised manner, significantly reducing labor-intensive annotation and the requirement of computational resources. 2) DCNN-based models ensure the classification task in a high accurate and computational efficiency. 3) the exploration and discovery utilities enable the discovery of the informative features in large-scale comparative analysis of WSI datasets.

**Figure 1.**
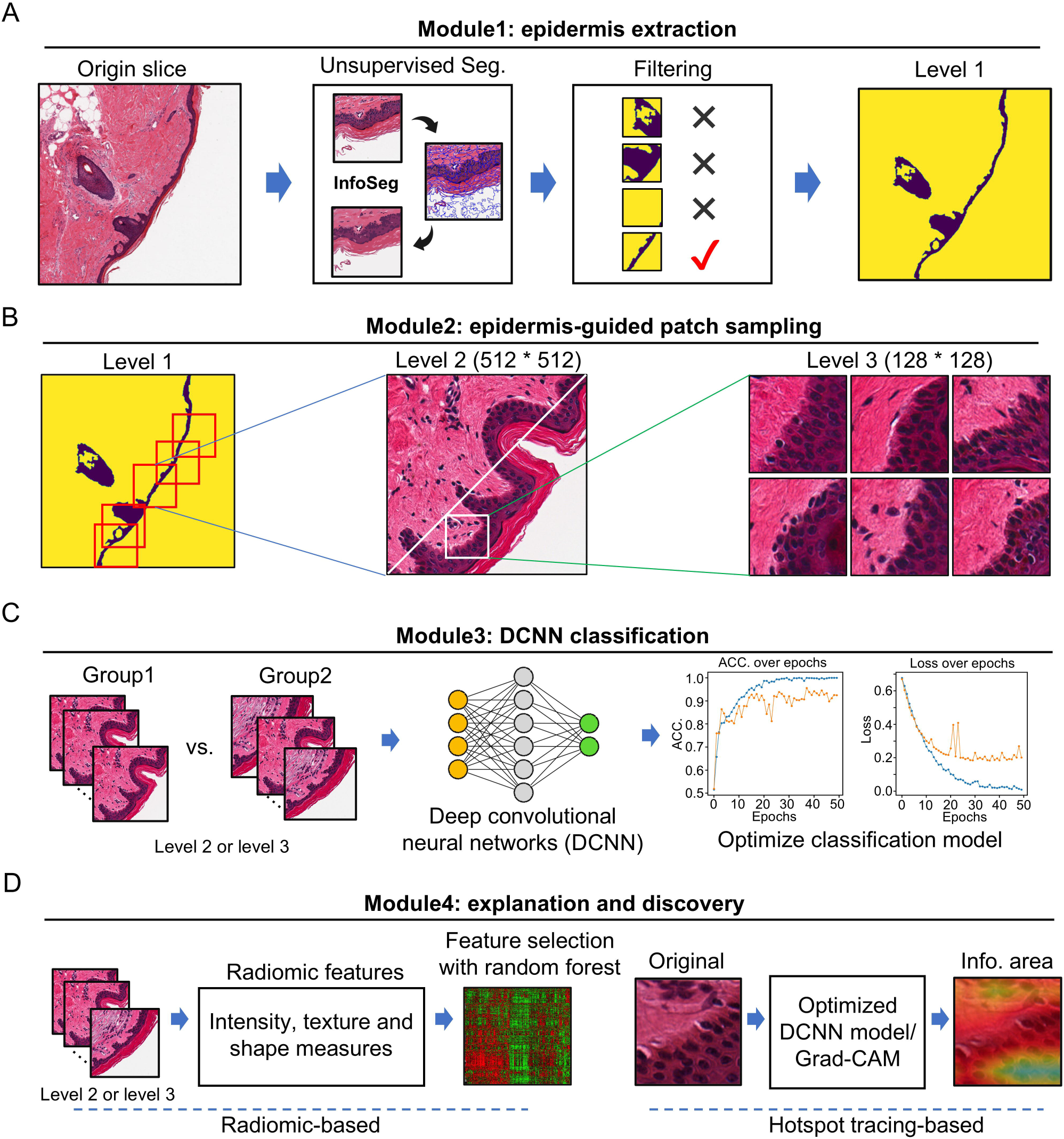
PathoEye: a deep learning framework for histopathological image analysis of skin tissue. A. the scheme for epidermis extraction in histopathological image of skin tissue; B. three levels of segregated images used in PathoEye analyses; C. DCNN classification model with level 2 image as input. D. the explanation and discovery module for WSI analysis.

### PathoEye outperforms the existing methods in classification tasks

PathoEye incorporates an innovative epidermis-guided patch sampling strategy to utilize the pathological alternation of BMZ in both level 2 and 3 images, which helps discriminate multiple skin conditions. To test this, we collected H&E stained WSI images of skin biopsy from the young (n=83) and aged (n=83) skin from GTEx project ^**31**^ for comparison analysis. Then, we compared the performance of PathoEye with CLAM ^**38**^, transMIL ^**39**^ and DSMIL ^**40**^ for the classification tasks because all of them incorporate RestNet50, but they extract features using different strategies (Figure 2A). Firstly, we compared their performance in discriminating the aged and young skin in level 2 images (Figure 2B). As a result, the receiver operating characteristic (ROC) curve and the precision-recall (PR) curve showed that PathoEye has better performance (ROC = 0.99, AUC = 0.99) compared with CLAM (ROC = 0.89, AUC = 0.92), TransMIL (ROC = 0.73, AUC = 0.77) and DSMIL (ROC = 0.87, AUC = 0.90), indicating that the patch sampling method with biological implication outperforms the method with attention algorithm or while bag selection strategies (Figure 2C and 2D). The boxplot showed that PathoEye performed better than the existing methods in the 5-fold cross-validation analyses in terms of accuracy, ROC, and PR (Figure 2E-G), suggesting the robustness of PathoEye in discriminating the young and aged skin using WSIs. Further, we compared the performance of PathoEye with the existing methods in binary- and multi-classification of skin diseases, including bullous pemphigoid (18), lichen planus (18), cutaneous amyloidosis (18), atopic dermatitis (18) (Figure 2H). The results showed that PathoEye has the same performance as CLAM and TransMIL in the binary classification tasks, while it has a better performance in the multi-classification tasks (Accuracy = 0.817, F1 = 0.729 and AUC = 0.961) (Figure 2I). Our analyses showed that the epidermis-guided patch sampling strategy improves the performance of the DCNN classification model, and PathoEye outperforms the existing methods in classification tasks.

**Figure 2.**
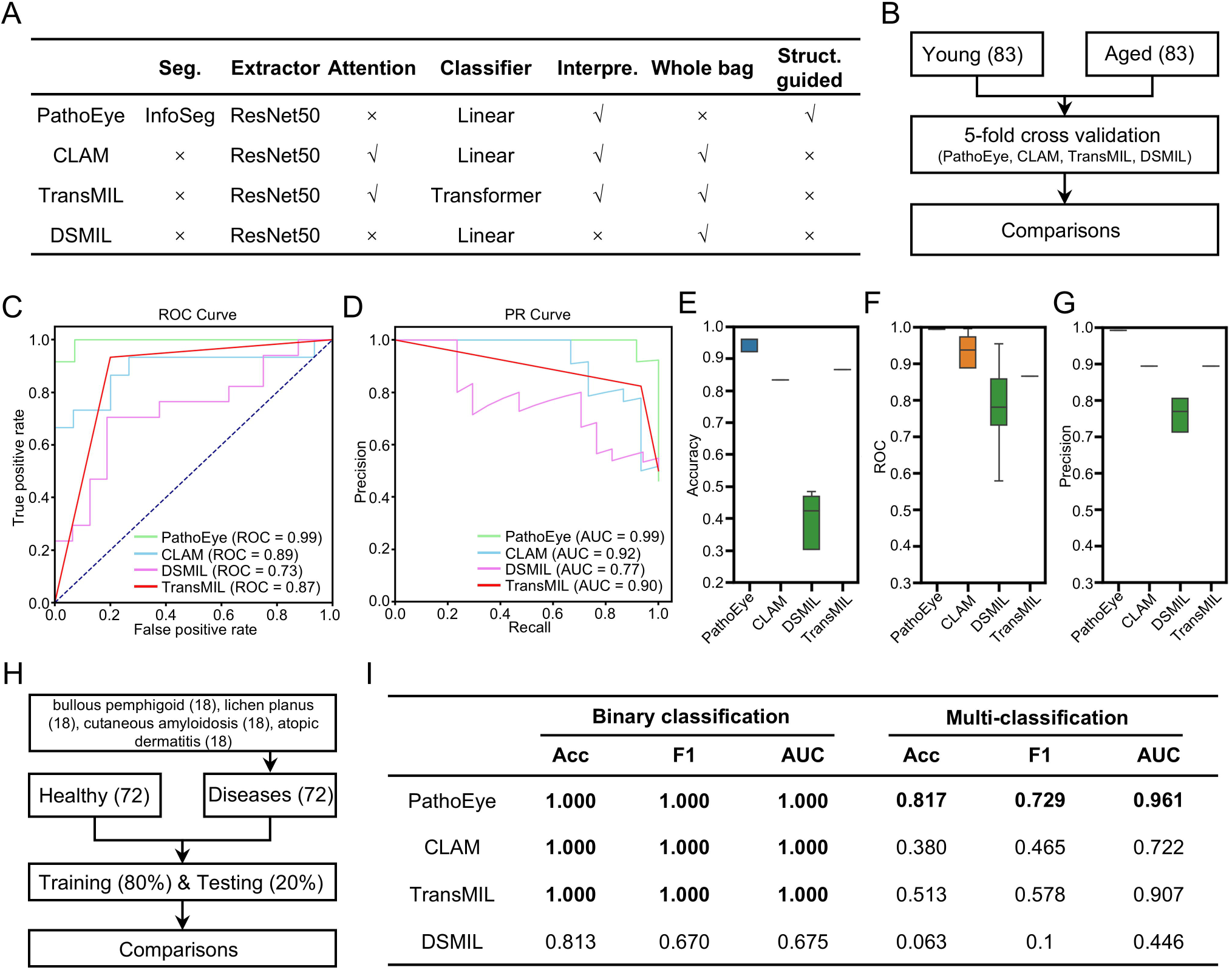
PathoEye outperforms the existing methods in classification tasks. A. the characteristics of PathoEye, CLAM, TransMIL and DSMIL in the comparison analyses. B. the analysis scheme of the binary classification of the young and aged skin tissues. C and D. the receiver operating characteristic (ROC) curve (C) and the precision-recall (PR) curve (D) showed that PathoEye has a better performance compared with CLAM, TransMIL or DSMIL. E-G. the boxplots showed that PathoEye performs better than the existing methods in the 5-fold cross-validation analyses in terms of accuracy (E), ROC (F), and PR (G). H. the scheme of classifying the diseased and the healthy skin tissues. I. the comparative analyses of the performance of 4 models.

### PathoEye shows a decrease in epidermis thickness and the variance of rete ridge length in the aged skin

Next, we sought to explore the features of H&E-stained WSIs of different ages using the exploration and discovery utilities in PathoEye. To this end, we collected a larger WSI dataset (n=917) of skin trusses derived from different ages and sun exposure conditions. Then, we calculated the epidermis rete ridge score as the variance of the shortest distance between the point of the basal layer to stratum corneum boundaries and the epidermis thickness as the mean distance between them (Figure 3A and Figure S3). As a result, we observed that the thickness of the epidermis was significantly decreased in aged skin in both sun-exposed and non-sun-exposed groups (Figure 3B and 3D, Table S2). Interestingly, we found that the epidermis thickness gradually decreased from 30 years old and remained stable from 60 years old (Figure 3B and 3D). In addition, we observed that the variance of rete ridge length of the epidermis-dermis boundary is gradually decreased during aging in both sun-exposed and non-sun-exposed groups (Figure 3C and 3E, Table S3). Notably, the decrease of variance of rete ridge length is initiated at 20 years old and remained constant at 50 years old, earlier than the decrease of epidermis thickness. Our analysis showed a strikingly decreased thickness and variance of rete ridge length of the epidermis during skin aging consistent with previous studies ^**41,42**^, demonstrating the utility of PathoEye in exploring quantitative features of WSIs.

**Figure 3.**
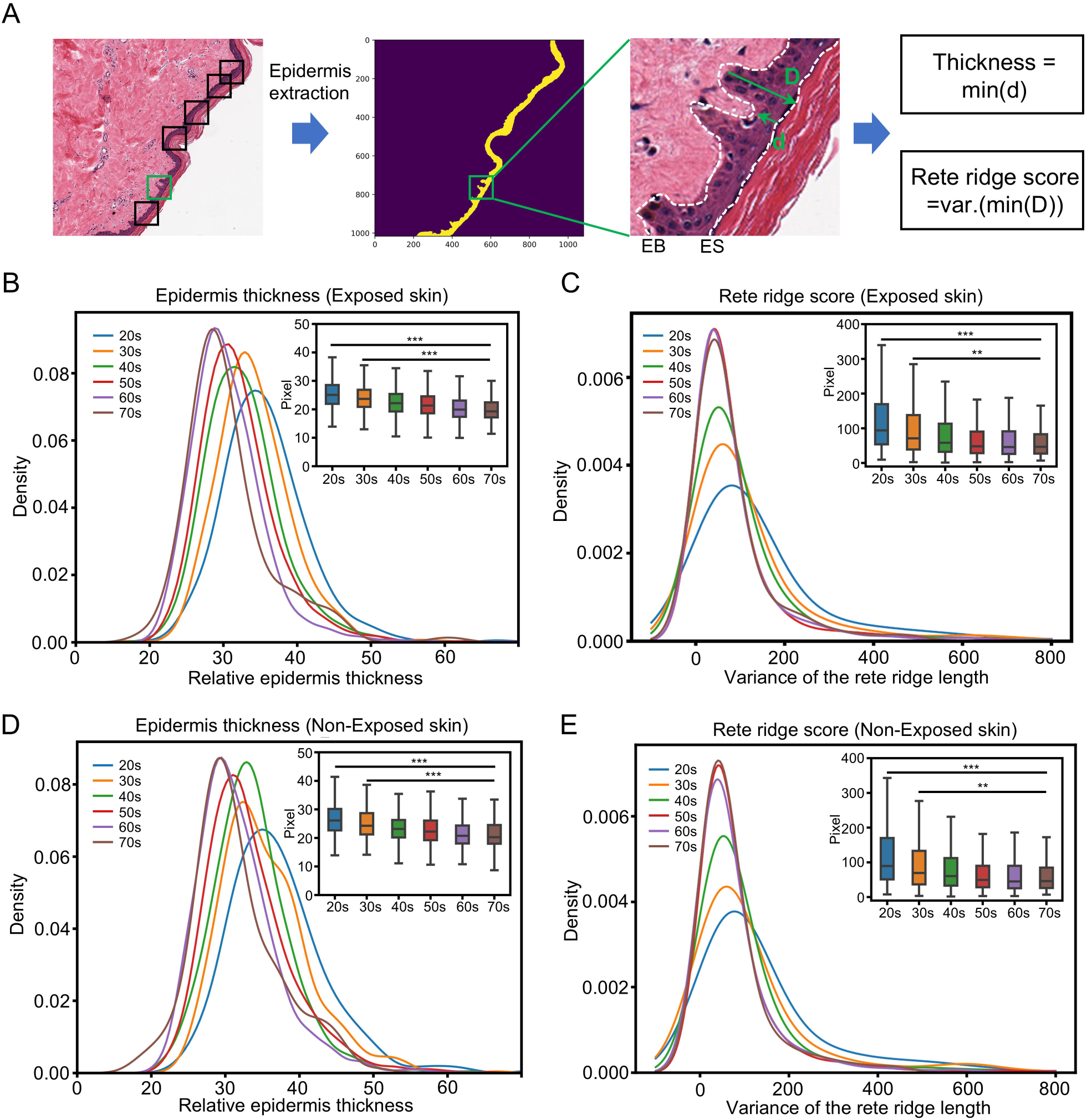
PathoEye shows a decrease in epidermis thickness and the variance of rete ridge length in the aged skin. A. the schematic diagram illustrates the definition of rete ridge score and thickness used to describe the morphology of skin epidermis. B. the distribution of epidermic thickness in skin tissue of different age stages in sun-exposed condition; C. the changes of rete ridge length of BM in skin tissue of different age stages in sun-exposed condition; D. the distribution of epidermic thickness in skin tissue of different age stages in non-sun-exposed condition; E. the changes of rete ridge length of the BMZ in skin tissue of different ages in non-sun-exposed condition. These figures demonstrated that the thickness and rete ridge length of the epidermis decreased along with skin aging.

### PathoEye identifies the aged-associated patterns in the epidermal layer and basement membrane zone

Then, we applied PathoEye to explore the informative features associated with the aged skin using the level 2 images derived from the young (20-29 years old) and aged (70-79 years old) skins. To avoid the influence of skin morphology, we only included the level 2 image containing BMZ (the dermis, epidermis, and epidermis diagonally across the patch). For the radiomic-based exploration, 101 radiomic features were subjected to clustering and prioritization analysis (Table S5), resulting in the top 5 features associated with the aged skin using the random forest (RF) algorithm (Figure 4A and 4B, Figure S4 and S5). Notably, the three most significant features are the first-order 10th percentile, the first-order median, and the Gray-Level Co-occurrence Matrix (GLCM) cluster shade, which is associated with the saturation of color and texture complexity of an image (Figure 4C). Expectedly, these features gradually increase or decrease along aging (Figure 4D-F) and the changes are indeed observed in epidermis (Figure 4G). For the hotspot tracing-based exploration, we traced the regions representing the differences between the young and aged skin using the established DCNN-based classification model (Figure S4, the bottom panel). As a result, we found that the epidermis region and BMZ are informative (the hotspot region) for discriminating between young skin and aged skin (Figure 4H and 4I). Our further investigation showed that the chronic aging-related effects in BMZ of the aged skin using a similar analysis scheme (Figure 4J and 4K, Figure S5A-C, Table S6 and Table S7), consistent with that the aberrations of BMZ are critical for epidermal stem cells (EpiSCs) functional decline in aged skin ^**20,43**^. In summary, PathoEye is a comprehensive tool for exploring the representative changes of different skin conditions in a zoom-in manner.

**Figure 4.**
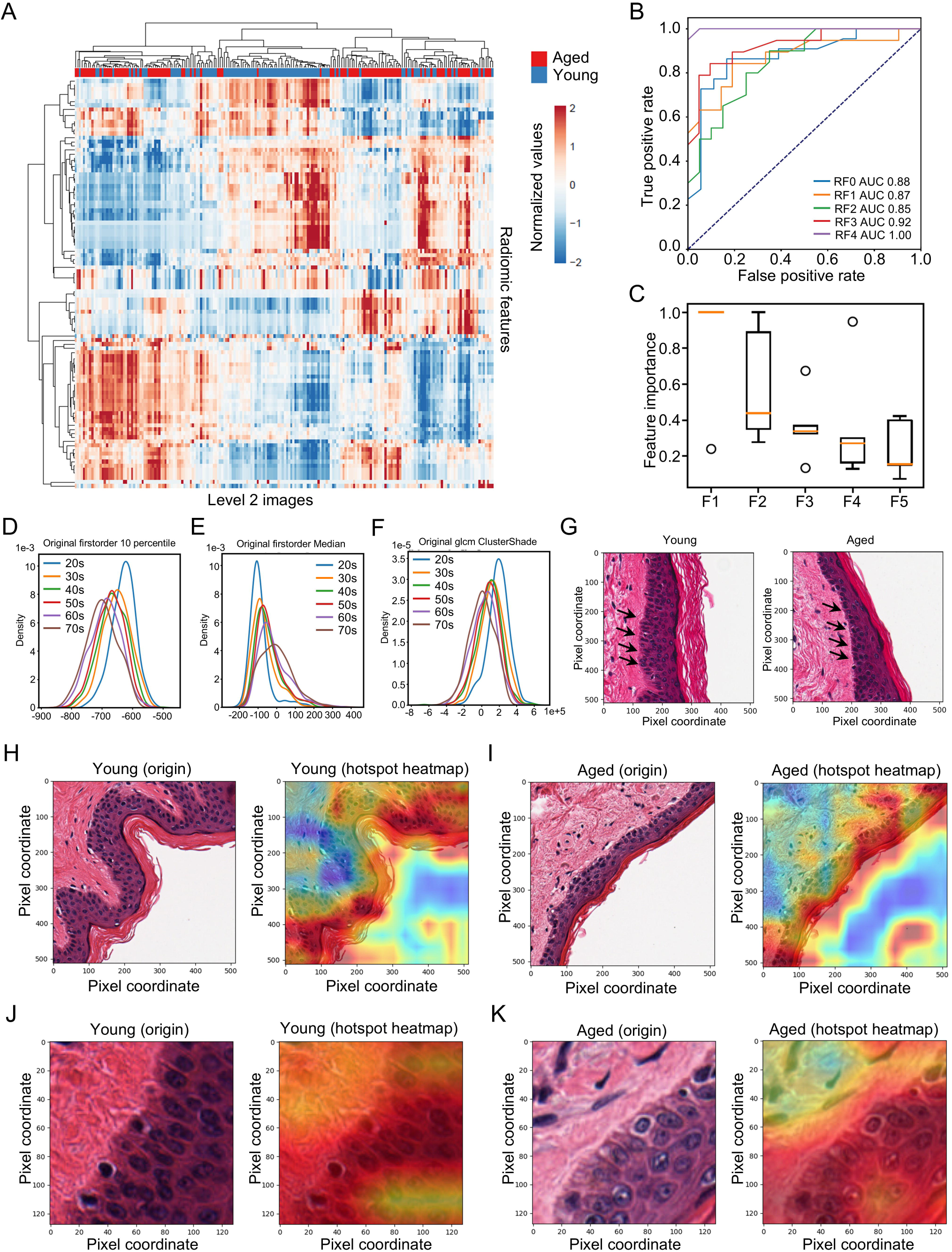
PathoEye identifies the aged-associated patterns in the epidermal layer and the basement membrane zone. A. the heatmap clustering analysis of the radiomic features extracted from level 2 images, indicating that the radiomic features are distinct between young and old skin; B. Receiver Operating Characteristic (ROC) curve and Area Under Curve (AUC) analyses of Random Forest model for discriminating the young and aged skin; C. the top 5 crucial features in Random Forest model; F1: original first order 10 Percentile; F2: original first order Median; F3: original glcm Cluster Shade; F4: original first order Skewness; F5: diagnostics image-original Mean. D-F. density plots show the distribution of original first order 10 Percentile (D), original first order Median (E) and original glcm Cluster Shade (F) along skin aging. G. a snapshot illustrates the observed defects in the young and aged skin tissue. H and I. the representative diagram showed the hotspot tracking of the level 2 image using Grad-CAM. J and K. the representative diagram showed the hotspot tracking of the level 3 image using Grad-CAM.

### Validation of the aged-associated defects observed in the aged skin

Next, we plan to validate the observed age-related defects experimentally. Our immunofluorescence (IF) staining revealed that EpiSCs located in the basement membrane (BM) have a higher expression of p16^INK4a^ in aged skin compared to young skin, indicating a higher percentage of senescent EpiSCs in aged skin (Figures 5A and 5B). Next, we investigated the protein expression of key components and regulators associated with the homeostasis of the BM and attached stem cells, including COL17A1, COL4A2, ITGA6, and PLEC ^**41,44–46**^. As expected, our immunohistochemistry (IHC) experiments showed elevated levels of COL17A1, COL4A2, ITGA6 and PLEC proteins in aged skin compared to young skin (see Figures 5C-J and Figure S6), suggesting a transition in skin stiffness from young to aged. In summary, our experimental findings confirm that more senescent EpiSCs reside in the BM of aged skin, along with defects in the membrane zone (BMZ) components, thereby underscoring the potential application of PathoEye in dermatological research.

**Figure 5.**
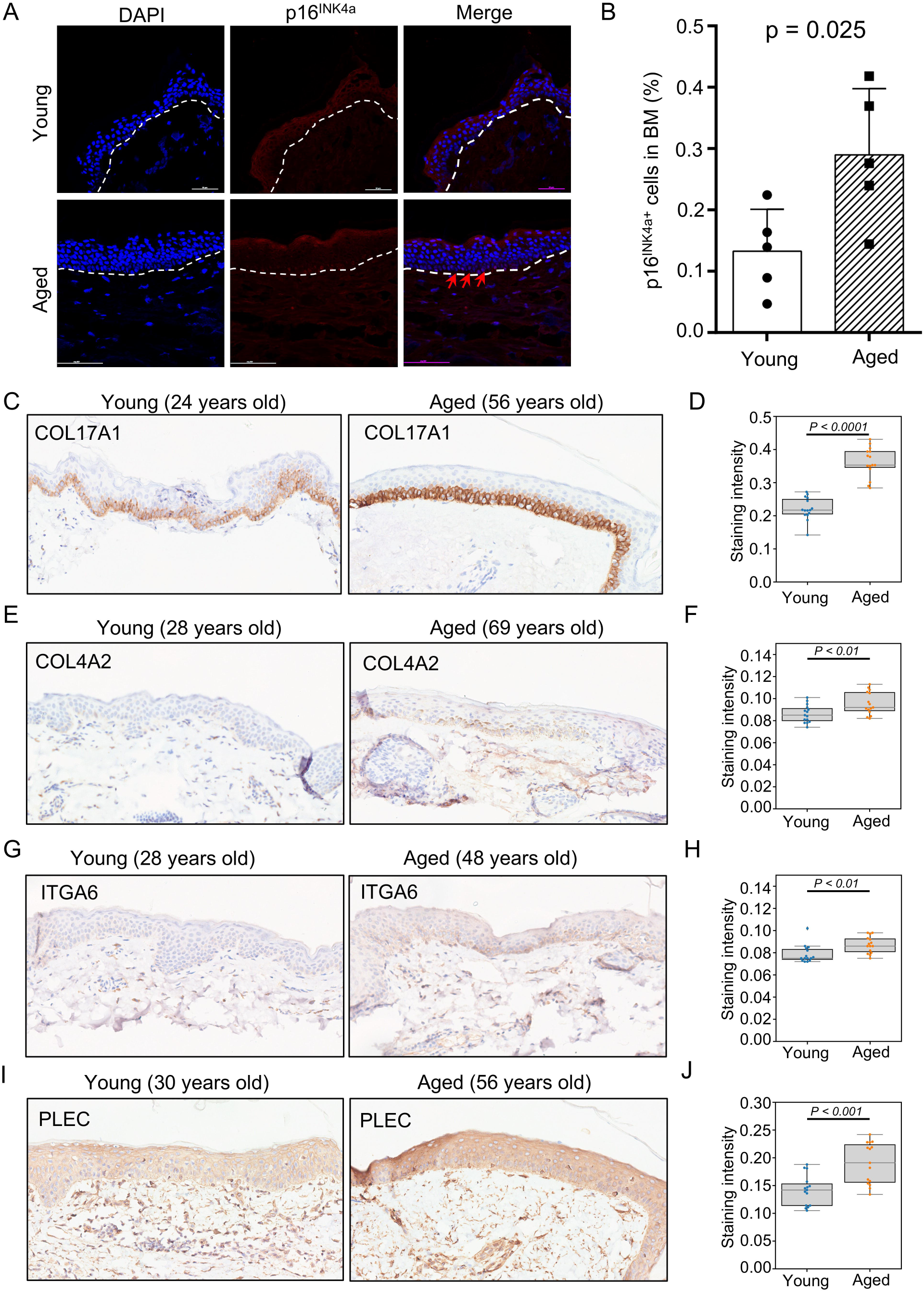
Validation of the aged-associated defects in aged skin. A. IF staining shows the aged EpiSCs residing along the BM in aged skin; B. the boxplot shows the dysregulated expression level of p16^INK4a^ in young and aged skin. C-J. IHC staining of COL17A1 (C and D), COL4A2 (E and F), ITGA6 (G and H) and PLEC (I and J) in the young and aged skin.

## Discussions

In this study, we demonstrated that PathoEye is an interpretable histological image analysis pipeline capable of extracting and exploring novel features of skin tissue with a training process with a few WSIs. PathoEye addresses several critical challenges in WSI analysis: its heavy reliance on manual annotation, the substantial computational resources required, the need for interpretability, and issues with generalization. It employs an unsupervised semantic segmentation method and an epidermis-guided patch sampling technique to reduce the need for labor-intensive manual annotation and enhance performance in classification tasks. Furthermore, PathoEye identifies the features related to skin aging in the basement membrane zone (BMZ). In the future, we aim to expand the applications of PathoEye to skin disease diagnosis and research.

The high resolution of WSIs and the multiple morphological layers of skin tissues pose significant challenges in the comparative analysis of large-scale datasets ^**22,47**^. Numerous studies have demonstrated the successful extraction of tumor entities, distinguishing skin tumors from normal skin samples using deep learning algorithms with WSIs as input ^**22,28,47,48**^. Recent research has indicated that the skin epidermis and BMZ provide valuable information for classification tasks, as most cytological features can be observed in the BMZ ^**41**^. PathoEye automatically unsupervised extracts the epidermis and downsamples the WSIs through a hierarchical strategy. The sampled patches contain the dermis, epidermis, and BMZ along the same diagonal line, representing most cytological features in skin biopsies. Our approach generates limited anatomically consistent images for radiomic feature extraction, category classification, and hotspot tracing in downstream analyses. This method eliminates the need for labor-intensive manual annotations and significantly reduces the computational resources required during the training and prediction process. With these advantages, our strategy enhances the application of computational pathology in diagnosing skin diseases and serves as a representative model for WSI analysis in dermatology.

Previous reports showed that many deep-learning models had achieved high accuracy in predicting melanoma or basal cell carcinoma ^**17,28,47**^. In practice, those models output the prediction label with WSI as input without any explainable evidence corresponding to their decisions ^**27**^. A few studies demonstrated that semantic segmentation and annotation of the skin structure help inform the morphological abnormalities in WSI analysis^**18,28**^. Here, we employed radiomic features and a DCNN-based hotspot tracing model to provide evidence supporting the decision of the classification tasks. The radiomic features represent the global texture of the down-sampled images, while the hotspot tracing focuses on the aberrations of different layers in skin tissue spatially. Our design ensures the application of PathoEye in clinical diagnosis and dermatological research.

## Code and data availability

The source code is available at github (https://github.com/lysovosyl/PathoEye), and the data is available at Zenodo (https://zenodo.org/records/10681896). CLAM, TransMIL and DSMIL were downloaded from github.

## Contributions

J.Z., Y.L. and B.Y. designed the project; Y.L. implemented the PathoEye algorithm and drafting the initial manuscript; Y.L., H.S., T.Y. and X.Z. performed data analyses; F.L., Y.Z. and H.J. collected clinical samples and performed IF/IHC analyses; J.W. and G.L. collected the H&E staining WSIs; J.Z., and Y.L. revised the manuscript; J.Z., Y.L, B.Y., H.S. and R.X. contributed to the conceptualization of the methods used and reviewed the draft. All authors read and approved the final manuscript.

## Reference

1. Alom MZ, Yakopcic C, Nasrin MstS, Taha TM, Asari VK. Breast Cancer Classification from Histopathological Images with Inception Recurrent Residual Convolutional Neural Network. J Digit Imaging 2019;32:605–17doi:10.1007/s10278-019-00182-7.

2. Bera K, Schalper KA, Rimm DL, Velcheti V, Madabhushi A. Artificial intelligence in digital pathology — new tools for diagnosis and precision oncology. Nat Rev Clin Oncol 2019;16:703–15.

3. Wong ANN, He Z, Leung KL, et al. Current Developments of Artificial Intelligence in Digital Pathology and Its Future Clinical Applications in Gastrointestinal Cancers. Cancers (Basel) 2022;14:3780doi:10.3390/cancers14153780.

4. Sakamoto T, Furukawa T, Lami K, et al. A narrative review of digital pathology and artificial intelligence: focusing on lung cancer. Transl Lung Cancer Res 2020;9:2255– 76doi:10.21037/tlcr-20-591.

5. Funkhouser WK. Pathology: The Clinical Description of Human Disease. Molecular Pathology 2009;:197–207doi:10.1016/B978-0-12-374419-7.00011-1.

6. Bueno G, Déniz O, Fernández-Carrobles MDM, Vállez N, Salido J. An automated system for whole microscopic image acquisition and analysis. Microsc Res Tech 2014;77:697–713.

7. Kumar N, Busarla SVP, Sayed S, et al. Telecytology in East Africa: a feasibility study of forty cases using a static imaging system. J Telemed Telecare 2012;18:7–12.

8. Jahn SW, Plass M, Moinfar F. Digital Pathology: Advantages, Limitations and Emerging Perspectives. J Clin Med 2020;9:3697doi:10.3390/jcm9113697.

9. Atabansi CC, Nie J, Liu H, Song Q, Yan L, Zhou X. A survey of Transformer applications for histopathological image analysis: New developments and future directions. BioMedical Engineering OnLine 2023;22:96doi:10.1186/s12938-023-01157-0.

10. Al-Thelaya K, Gilal NU, Alzubaidi M, et al. Applications of discriminative and deep learning feature extraction methods for whole slide image analysis: A survey. Journal of Pathology Informatics 2023;14:100335doi:10.1016/j.jpi.2023.100335.

11. Niazi MKK, Parwani AV, Gurcan MN. Digital pathology and artificial intelligence. The Lancet Oncology 2019;20:e253–61doi:10.1016/S1470-2045(19)30154-8.

12. Yoshikawa-Murakami C, Mizutani Y, Ryu A, et al. A Novel Method for Visualizing Melanosome and Melanin Distribution in Human Skin Tissues. Int J Mol Sci 2020;21:8514.

13. Nofallah S, Mokhtari M, Wu W, et al. Segmenting Skin Biopsy Images with Coarse and Sparse Annotations using U-Net. J Digit Imaging 2022;35:1238– 49doi:10.1007/s10278-022-00641-8.

14. Wei L, Gan Q, Ji T. Skin Disease Recognition Method Based on Image Color and Texture Features. Comput Math Methods Med 2018;2018:8145713doi:10.1155/2018/8145713.

15. Goyal M, Knackstedt T, Yan S, Hassanpour S. Artificial intelligence-based image classification methods for diagnosis of skin cancer: Challenges and opportunities. Computers in Biology and Medicine 2020;127:104065doi:10.1016/j.compbiomed.2020.104065.

16. Otaka H, Shimakura H, Motoyoshi I. Perception of human skin conditions and image statistics. J Opt Soc Am A, JOSAA 2019;36:1609–16doi:10.1364/JOSAA.36.001609.

17. Zheng T, Chen W, Li S, et al. Learning how to detect: A deep reinforcement learning method for whole-slide melanoma histopathology images. Comput Med Imaging Graph 2023;108:102275.

18. Xu H, Lu C, Berendt R, Jha N, Mandal M. Automated analysis and classification of melanocytic tumor on skin whole slide images. Comput Med Imaging Graph 2018;66:124–34.

19. Lu C, Mandal M. Automated analysis and diagnosis of skin melanoma on whole slide histopathological images. Pattern Recognition 2015;48:2738– 50doi:10.1016/j.patcog.2015.02.023.

20. Sekiguchi R, Yamada KM. Basement membranes in development and disease. Curr Top Dev Biol 2018;130:143–91doi:10.1016/bs.ctdb.2018.02.005.

21. Tsutsui K, Machida H, Nakagawa A, et al. Mapping the molecular and structural specialization of the skin basement membrane for inter-tissue interactions. Nat Commun 2021;12:2577doi:10.1038/s41467-021-22881-y.

22. Chen RJ, Ding T, Lu MY, et al. Towards a general-purpose foundation model for computational pathology. Nat Med 2024;:1–13doi:10.1038/s41591-024-02857-3.

23. Majumder S, Katz S, Kontos D, Roshkovan L. State of the art: radiomics and radiomics-related artificial intelligence on the road to clinical translation. BJR Open 2024;6:tzad004.

24. Gillies RJ, Kinahan PE, Hricak H. Radiomics: Images Are More than Pictures, They Are Data. Radiology 2016;278:563–77doi:10.1148/radiol.2015151169.

25. Lambin P, Rios-Velazquez E, Leijenaar R, et al. Radiomics: Extracting more information from medical images using advanced feature analysis. Eur J Cancer 2012;48:441–6doi:10.1016/j.ejca.2011.11.036.

26. McCague C, Ramlee S, Reinius M, et al. Introduction to radiomics for a clinical audience. Clinical Radiology 2023;78:83–98doi:10.1016/j.crad.2022.08.149.

27. Aeffner F, Zarella MD, Buchbinder N, et al. Introduction to Digital Image Analysis in Whole-slide Imaging: A White Paper from the Digital Pathology Association. Journal of Pathology Informatics 2019;10:9doi:10.4103/jpi.jpi_82_18.

28. Wang L, Shao A, Huang F, et al. Deep learning-based semantic segmentation of non-melanocytic skin tumors in whole-slide histopathological images. Experimental Dermatology 2023;32:831–9doi:10.1111/exd.14782.

29. Xu H, Usuyama N, Bagga J, et al. A whole-slide foundation model for digital pathology from real-world data. Nature 2024;630:181–8.

30. Selvaraju RR, Cogswell M, Das A, Vedantam R, Parikh D, Batra D. Grad-CAM: Visual Explanations from Deep Networks via Gradient-based Localization. Int J Comput Vis 2020;128:336–59doi:10.1007/s11263-019-01228-7.

31. GTEx Consortium. The Genotype-Tissue Expression (GTEx) project. Nat Genet 2013;45:580–5.

32. van Griethuysen JJM, Fedorov A, Parmar C, et al. Computational Radiomics System to Decode the Radiographic Phenotype. Cancer Res 2017;77:e104–7.

33. He K, Zhang X, Ren S, Sun J. Deep Residual Learning for Image Recognition. 2015 doi:10.48550/arXiv.1512.03385.

34. Chen F, Zhou J, Li Y, et al. YY1 regulates skeletal muscle regeneration through controlling metabolic reprogramming of satellite cells. EMBO J 2019;38.

35. Pérez-Anker J, Ribero S, Yélamos O, et al. Basal cell carcinoma characterization using fusion ex vivo confocal microscopy: a promising change in conventional skin histopathology. Br J Dermatol 2020;182:468–76.

36. Tizhoosh HR, Pantanowitz L. Artificial Intelligence and Digital Pathology: Challenges and Opportunities. Journal of Pathology Informatics 2018;9:38doi:10.4103/jpi.jpi_53_18.

37. Harb R, Knöbelreiter P. InfoSeg: Unsupervised Semantic Image Segmentation with Mutual Information Maximization. 2021doi:10.48550/arXiv.2110.03477.

38. Lu MY, Williamson DFK, Chen TY, Chen RJ, Barbieri M, Mahmood F. Data-efficient and weakly supervised computational pathology on whole-slide images. Nat Biomed Eng 2021;5:555–70.

39. Shao Z, Bian H, Chen Y, et al. TransMIL: Transformer based Correlated Multiple Instance Learning for Whole Slide Image Classification. 2021doi:10.48550/arXiv.2106.00908.

40. Yan Y, Wang X, Guo X, Fang J, Liu W, Huang J. Deep Multi-instance Learning with Dynamic Pooling. 2018https://www.semanticscholar.org/paper/Deep-Multi-instance-Learning-with-Dynamic-Pooling-Yan-Wang/0c383b30869e1af2b587043bab88b096d52552be. Accessed 3 Sep 2024.

41. Liu N, Matsumura H, Kato T, et al. Stem cell competition orchestrates skin homeostasis and ageing. Nature 2019;568:344–50doi:10.1038/s41586-019-1085-7.

42. Koester J, Miroshnikova YA, Ghatak S, et al. Niche stiffening compromises hair follicle stem cell potential during ageing by reducing bivalent promoter accessibility. Nat Cell Biol 2021;23:771–81.

43. Aleemardani M, Trikić MZ, Green NH, Claeyssens F. The Importance of Mimicking Dermal-Epidermal Junction for Skin Tissue Engineering: A Review. Bioengineering (Basel) 2021;8:148.

44. Verbeek E, Meuwissen ME, Verheijen FW, et al. COL4A2 mutation associated with familial porencephaly and small-vessel disease. Eur J Hum Genet 2012;20:844– 51doi:10.1038/ejhg.2012.20.

45. Matsumura H, Liu N, Nanba D, et al. Distinct types of stem cell divisions determine organ regeneration and aging in hair follicles. Nat Aging 2021;1:190– 204doi:10.1038/s43587-021-00033-7.

46. Yin H, Han S, Cui C, Wang Y, Li D, Zhu Q. Plectin regulates Wnt signaling mediated-skeletal muscle development by interacting with Dishevelled-2 and antagonizing autophagy. Gene 2021;783:145562.

47. Campanella G, Hanna MG, Geneslaw L, et al. Clinical-grade computational pathology using weakly supervised deep learning on whole slide images. Nat Med 2019;25:1301–9.

48. Yacob F, Siarov J, Villiamsson K, et al. Weakly supervised detection and classification of basal cell carcinoma using graph-transformer on whole slide images. Sci Rep 2023;13:7555.

