## Supplemental Figure for "PathoEye: a deep learning framework for the whole-slide image analysis of skin tissue"

Figure S1.

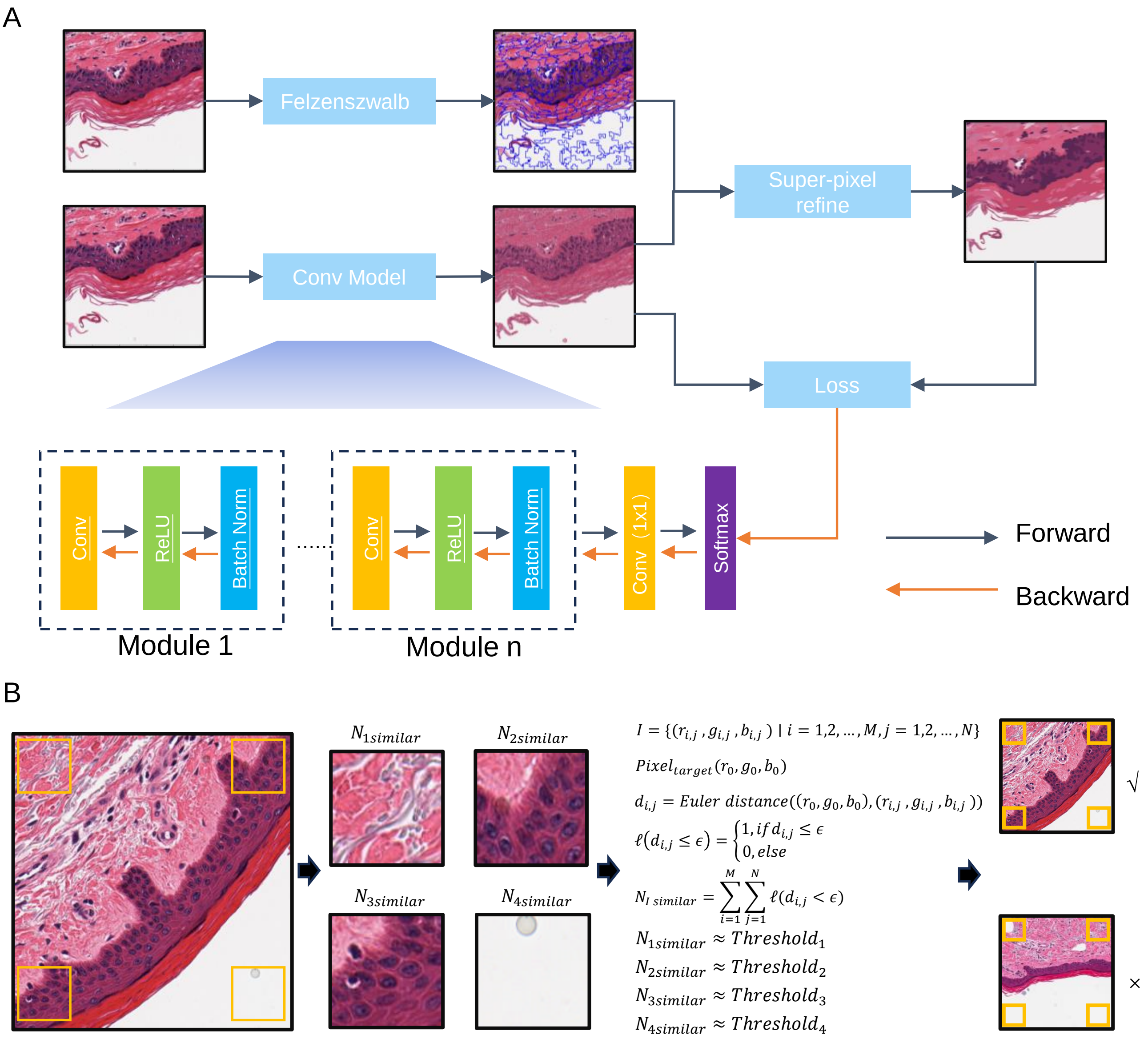


**Figure S1. The architecture of epidermis extraction (infoSeg model) and patch extraction with epidermis running diagonally.** A. it comprises a series of modules, including convolutional, ReLU, and batch normalization layers, including a 1x1 convolution layer and a softmax layer for pixel classification. Initially, an image is processed using the Felzenswalb algorithm to generate super-pixels. Then, the image is fed into the model to produce a mask, and the mask is refined based on the super-pixels, resulting in a new mask. The new mask will optimize the current model, enhancing its ability to generate improved masks in subsequent iterations. This process is repeated several times for each image until saturation. B. The scheme for extracting patches with epidermis running diagonally. Four small patches were sampled from each point of diagonal lines. For each patch, the similarity among patches was evaluated using *Euclidean distance*. Briefly, the higher similarity means the same pattern of two patches; then, the level 2 or level 3 images with epidermis diagonally were determined if $N_{3similar} \approx N_{2similar}$ and $N_{1similar}$ distinct from $N_{4similar}$.

Figure S2.

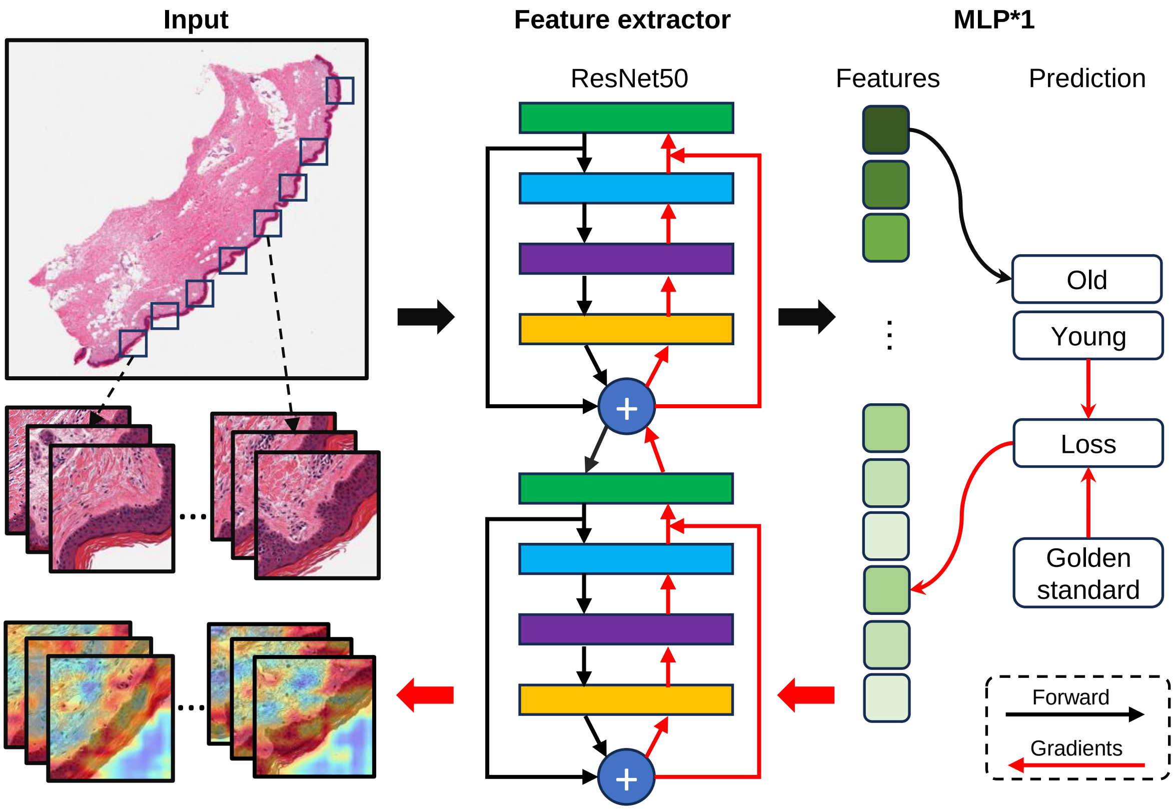


**Figure S2. The architecture of DCNN for discriminating the young and aged skin.** The DCNN model was trained to discriminated young and aged skin using the level 2 or level 3 images as input. Then, we traced the weighted aggregation of the feature maps and gradients in DCNN using the Grad-CAM algorithm to highlight the hotspot regions that were informative for prediction with the loss value, ground truth (golden standard), and prediction results.

Figure S3.

A

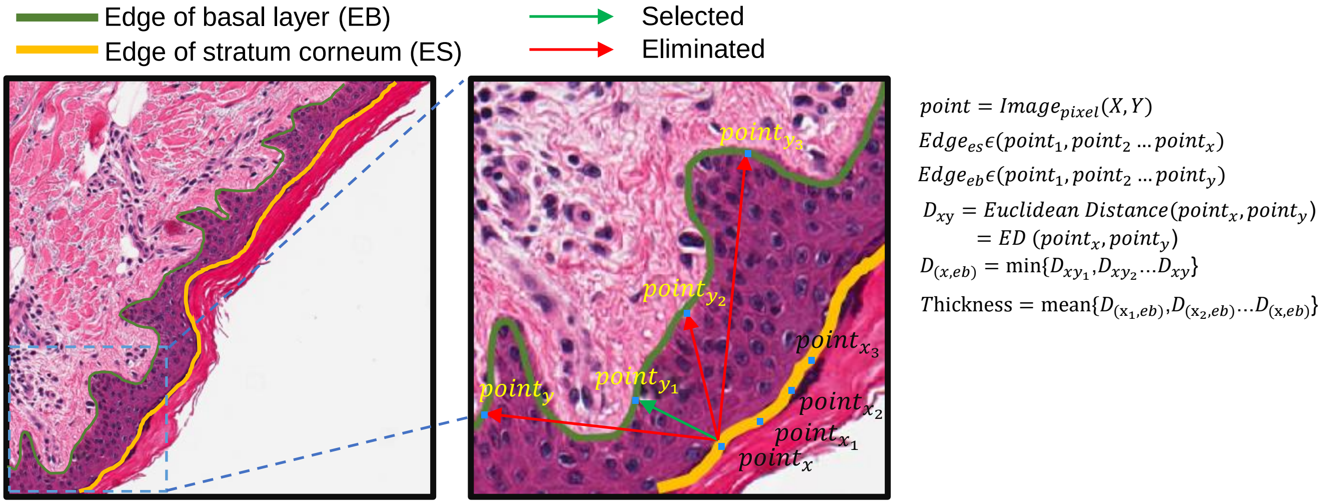


B

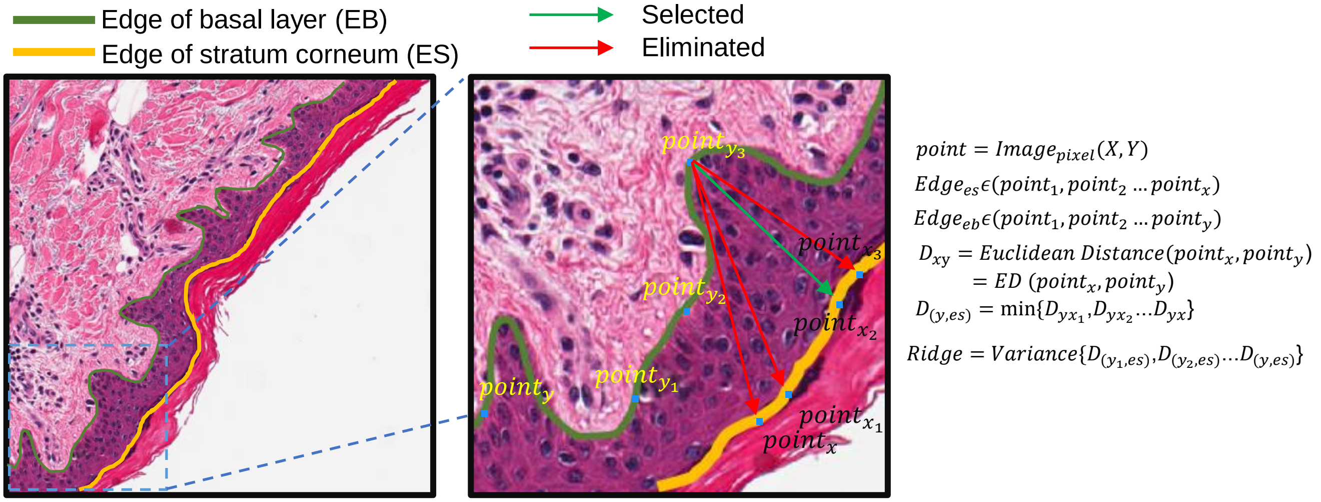


**Figure S3. The scheme for the thickness and the rete ridge length calculation of the epidermis.** A. The illustration represents the formula and calculation method for the thickness of the epidermis. We defined the mean of the shortest distant between piont$_x$ to point$_y$ as the thickness of epidermis. B. The illustration represents the formula and calculation method for the rete ridge length variation of epidermis. We defined the rete ridge length as the points of basal layer as the shortest distant from the edge of basal layer to the edge of stratum corneum. Therefore, the variation of the rete ridge length along the edge of the basal layer represents the degree of fold in the epidermis.

Figure S4.

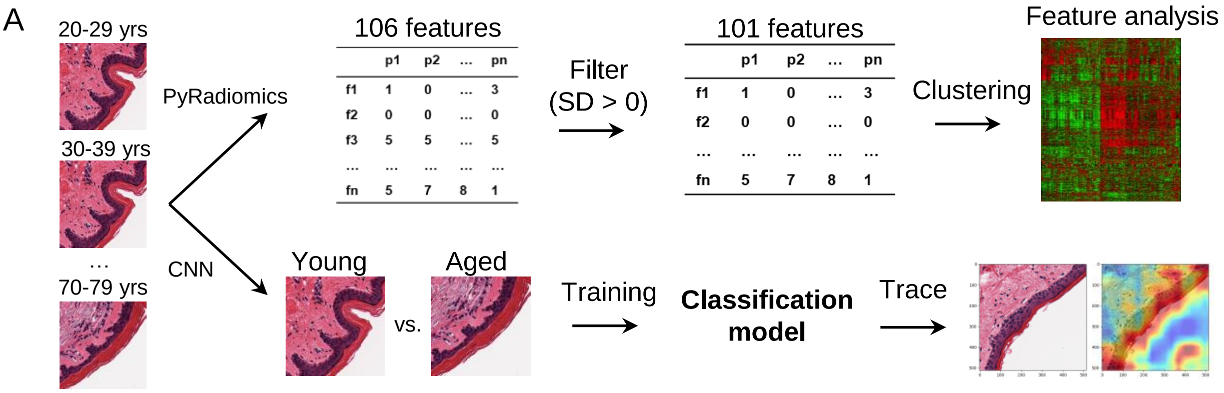


**Figure S4. The flowchart of WSI analysis derived from skin tissues of different age ranges.** The automated segmented images of varying age ranges were subjected to 1) radiomic feature analysis using PyRadiomics (top panel) and 2) classification analysis using CNN models (bottom panel).

Figure S5.

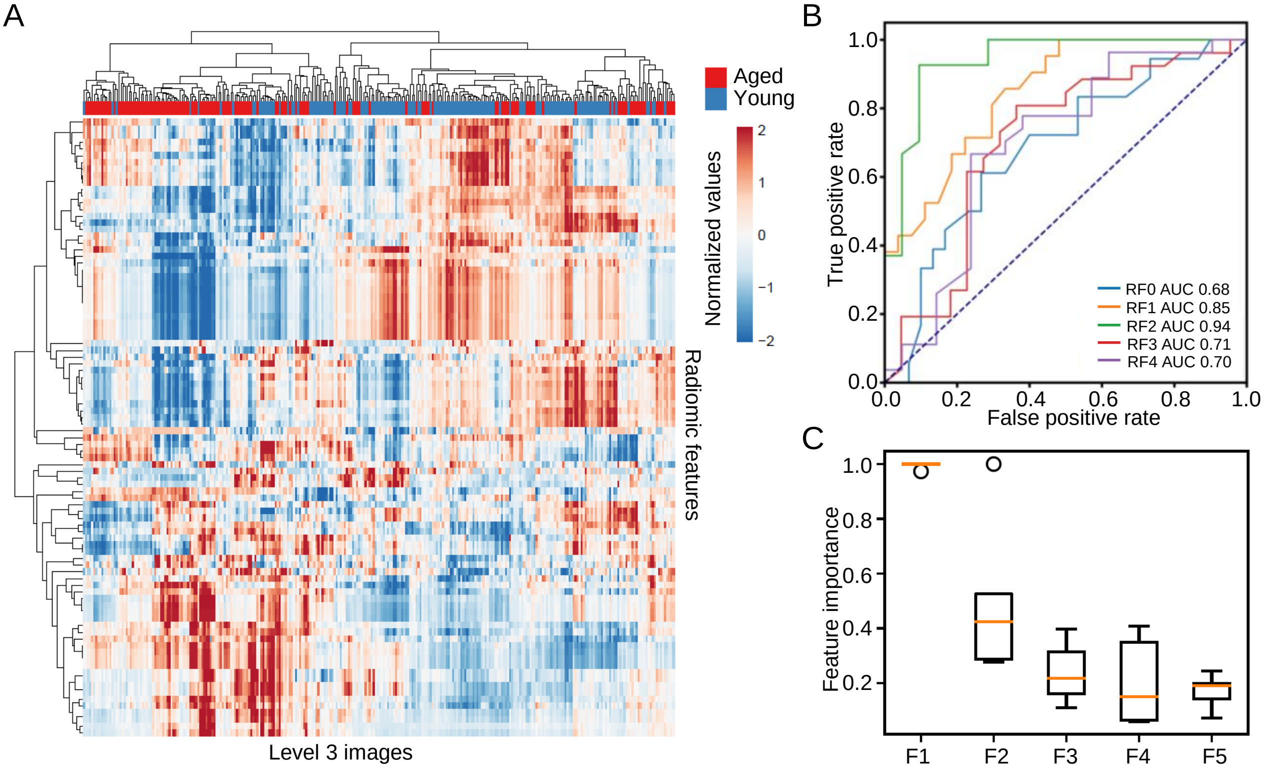


**Figure S5. PathoEye discovers the defects of the basement membrane zone in aged skin.** A. the heatmap cluster analysis of the radiomic features extracted from the level 3 images; B. Receiver Operating Characteristic (ROC) curve and Area Under Curve (AUC) analyses of the Random Forest model for discriminating the young and aged skin; C. the top 5 important features from the Random Forest model; F1: original first order 10 Percentile; F2: original first order Median; F3: original glszm GrayLevelVariance; F4: original glcm DifferenceVariance; F5: original glcm MCC.

Figure S6.

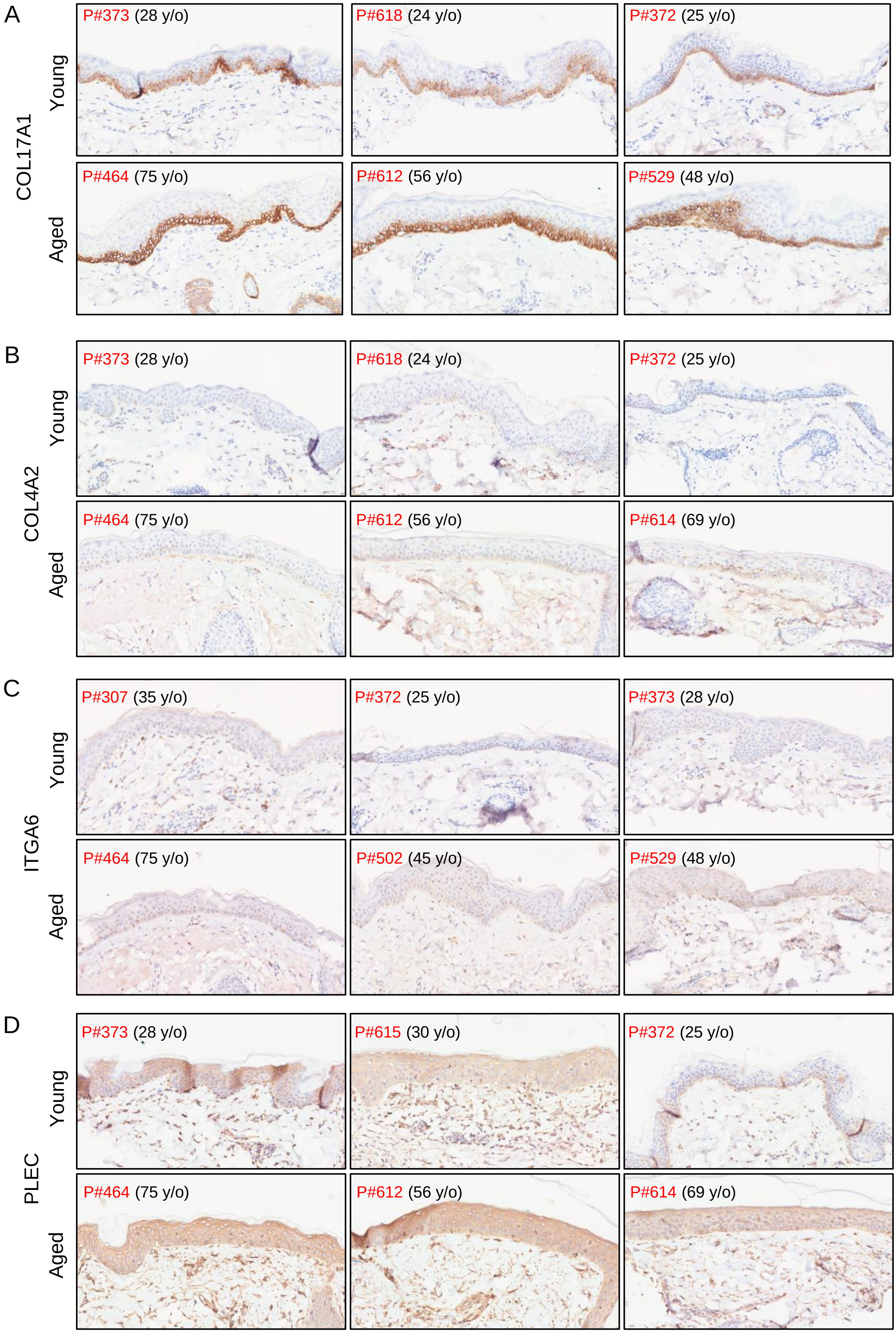


**Figure S6. The expression level of four extracellular genes in young and aged human skin.** Immunohistochemistry (IHC) staining for COL17A1 (A), COL4A2 (B), ITGA6 (C) and PLEC (D).
