## Supplemental Note for "PathoEye: a deep learning framework for the whole-slide image analysis of skin tissue"

Supplementary Notes

Epidermis extraction

We adopted the InfoSeg ^39^ algorithm to establish the epidermis extraction module, which leverages the benefit of the unsupervised segmentation deep learning algorithm and a customized image filter strategy (Figure 1A). Briefly, it is composited with three parts: 1) the pixels with similar patterns were clustered in a superpixel using the Felzenszwalb algorithm ^40^, those superpixels will be used to guide the optimization of the deep learning process; 2) a chain of deep learning modules consists of a convolutional layer, a rectified linear unit (ReLU) activation layer, and a batch normalization layer. Then, the model employs a 1 x 1 convolutional layer followed by a Softmax layer for pixel classification, resulting in a mask representing the epidermis region after several iterations (Figure S1A); 3) a customized image filtering process was employed to adjust the final mask using a corrosion expansion algorithm and a perfect mask as level 1 image of BMZ. The image and the informative mask were subjected to downstream analyses.

Epidermis-guided patch sampling

A customized pipeline was established to simulate the zoom-in and anatomical inspection process in WSI analysis (Figure 1B). Briefly, it generates 512 x 512-pixel patches (term as level 2 images) along the mask representing the epidermis region in WSIs, which composite with the dermis, BM, epidermis, and stratum corneum; then, 128 x 128-pixel patches (term as level 3 images) were extracted from the level 2 images along the mask representing epidermis; they represent the detailed information of BMZ. Finally, the level 2 and 3 images are retained for downstream analyses using a customized filtering method to ensure the BM or epidermis diagonally runs across the patch (Figure 1B and Figure S1B).

**Radiomic features calculation**

For each WSI of skin tissue, we calculated 106 features using Pyradiomics package, resulting in 18 First Order features, 24 Gray Level Co-occurrence Matrix features, 16 Gray Level Size Zone Matrix features, 16 Gray Level Run Length Matrix features, 5 Neighbouring Gray Tone Difference Matrix features, 14 Gray Level Dependence Matrix features and 13 diagnostics. The calculation of those features uses the equation listed as follows:

**First Order Features**

1. **Energy**

$$energy=\sum_{i=1}^{N_{p}} {(X\left( i \right)+c)}^{2}$$

2. **Total Energy**

$$total energy=V_{voxel}\sum_{i=1}^{N_{p}} {(X\left( i \right)+c)}^{2}$$

3. **Entropy**

$$entropy=-\sum_{i=1}^{N_{g}} p(i){log}_{2}(p\left( i \right)+\epsilon)$$

$\epsilon$is a arbitrarily small positive number $(\approx2.2\times{10}^{-16})$

4. **Minimum**

$$minimum=min(X)$$

5. 10th percentile

The 10th percentile of X

6. 90th percentile

The 90th percentile of X

7. Maximum

$$maximum=max(X)$$

8. Mean

$$mean=\frac{1}{N_{p}}\sum_{i=1}^{N_{p}} X(i)$$

9. Median

The median gray level intensity within the ROI.

10. Interquartile Range

$$interquartile range=P_{75}-P_{25}$$

11. Range

$$range=\max\left( X \right)-\min\left( X \right)$$

12. Mean Absolute Deviation (MAD)

$$MAD=\frac{1}{N_{p}}\sum_{i=1}^{N_{p}} |X\left( i \right)-\overline{X}|$$

13. Robust Mean Absolute Deviation (rMAD)

$$rMAD=\frac{1}{N_{10-90}}\sum_{i=1}^{N_{10-90}} |X_{10-90}\left( I \right)-\overline{X}_{10-90}|$$

14. Root Mean Squared (RMS)

$$RMS=\sqrt{\frac{1}{N_{p}}\sum_{i=1}^{N_{p}} {(X\left( i \right)+c)}^{2}}$$

15. Skewness

$$skewness=\frac{\mu^{3}}{\sigma^{3}}=\frac{\frac{1}{N_{p}}\sum_{i=1}^{N_{p}} {(X\left( i \right)-\overline{X})}^{3}}{\left( \sqrt{\frac{1}{N_{p}}\sum_{i=1}^{N_{p}} {(X\left( i \right)-\overline{X})}^{2}} \right)^{3}}$$

16. Kurtosis

$$kurtosis=\frac{\mu^{4}}{\sigma^{4}}=\frac{\frac{1}{N_{p}}\sum_{i=1}^{N_{p}} {(X\left( i \right)-\overline{X})}^{4}}{\left( \frac{1}{N_{p}}\sum_{i=1}^{N_{p}} {(X\left( i \right)-\overline{X})}^{2} \right)^{2}}$$

17. Variance

$$variance=\frac{1}{N_{p}}\sum_{i=1}^{N_{p}} {(X\left( i \right)-\overline{X})}^{2}$$

18. Uniformity

$$uniformity=\sum_{i=1}^{N_{g}} {p(i)}^{2}$$

**Gray Level Co-occurrence Matrix (GLCM) Features**

1. **Autocorrelation**

$$autocorrelation=\sum_{i=1}^{N_{g}} \sum_{j=1}^{N_{g}} p(i,j)ij$$

2. Joint Average

$$joint average=\sum_{i=1}^{N_{g}} \sum_{j=1}^{N_{g}} p(i,j)i$$

3. Cluster Prominence

$$cluster prominence=\sum_{i=1}^{N_{g}} \sum_{j=1}^{N_{g}} {(i+j-u_{x}-u_{y})}^{4}p(i,j)$$

4. Cluster Shade

$$cluster shade=\sum_{i=1}^{N_{g}} \sum_{j=1}^{N_{g}} {(i+j-u_{x}-u_{y})}^{3}p(i,j)$$

5. Cluster Tendency

$$cluster shade=\sum_{i=1}^{N_{g}} \sum_{j=1}^{N_{g}} {(i+j-u_{x}-u_{y})}^{2}p(i,j)$$

6. Contrast

$$contrast=\sum_{i=1}^{N_{g}} \sum_{j=1}^{N_{g}} {(i-j)}^{2}p(i,j)$$

7. Correlation

$$correlation=\frac{\sum_{i=1}^{N_{g}} \sum_{j=1}^{N_{g}} p(i,j)ij-\mu_{x}\mu_{y}}{\sigma_{x}(i)\sigma_{y}(j)}$$

8. Difference Average

$$difference average=\sum_{k=0}^{N_{g}-1} kp_{x-y}(k)$$

9. Difference Entropy

$$difference entropy=\sum_{k=0}^{N_{g}-1} p_{x-y}(k){log}_{2}(p_{x-y}\left( k \right)+\epsilon)$$

10. Difference Variance

$$difference variance=\sum_{k=0}^{N_{g}-1} {(k-DA)}^{2}p_{x-y}(k)$$

11. Joint Energy

$$joint energy=\sum_{i=1}^{N_{g}} \sum_{j=1}^{N_{g}} {(p(i,j))}^{2}$$

12. Joint Entropy

$$Joint Entropy=-\sum_{i=1}^{N_{g}} \sum_{j=1}^{N_{g}} p(i,j){log}_{2}(p\left( i,j \right)+\epsilon)$$

13. Informational Measure of Correlation (IMC) 1

$$IMC1=\frac{HXY-HXY1}{max\{HX,HY\}}$$

14. Informational Measure of Correlation (IMC) 2

$$IMC1=\sqrt{1-e^{-2(HXY2-HXY)}}$$

15. Inverse Difference Moment (IDM)

$$IDM=\sum_{k=0}^{N_{g}-1} \frac{p_{x-y}(k)}{1+k^{2}}$$

16. Maximal Correlation Coefficient (MCC)

$$MCC=\sqrt{second largest eigenvalue\left( \sum_{k=0}^{N_{g}} \frac{p\left( i,k \right)p(j,k)}{p_{x}(i)p_{y}(k)} \right)}$$

17. Inverse Difference Moment Normalized (IDMN)

$$IDMN=\sum_{k=0}^{N_{g-1}} \frac{p_{x-y}(k)}{1+\left( \frac{k^{2}}{N_{g}^{2}} \right)}$$

18. Inverse Difference (ID)

$$ID=\sum_{k=0}^{N_{g}-1} \frac{p_{x-y}(k)}{1+k}$$

19. Inverse Difference Normalized (IDN)

$$IDN=\sum_{k=0}^{N_{g}-1} \frac{p_{x-y}(k)}{1+\left( \frac{k}{N_{g}} \right)}$$

20. Inverse Variance

$$inverse variance=\sum_{k=1}^{N_{g}-1} \frac{p_{x-y}(k)}{k^{2}}$$

21. Maximum Probability

$$Maximum Probability=max(p(i,j))$$

22. Sum Average

$$Sum Average=\sum_{k=2}^{{2N}_{g}} p_{x+y}(k)k$$

23. Sum Entropy

$$Sum Entropy=\sum_{k=2}^{{2N}_{g}} p_{x+y}(k)\log_{2} (p_{x+y}\left( k \right)+\epsilon)$$

24. Sum of Squares

$$Sum of Squares=\sum_{i=1}^{N_{g}} \sum_{j=1}^{N_{g}} \left( i-\mu_{x} \right)^{2}p\left( i,j \right)$$

**Gray Level Size Zone Matrix (GLSZM) Features**

1. Small Area Emphasis (SAE)

$$SAE=\frac{\sum_{i=1}^{N_{g}} \sum_{j=1}^{N_{s}} \frac{P(i,j)}{j^{2}}}{N_{z}}$$

2. Large Area Emphasis (LAE)

$$LAE=\frac{\sum_{i=1}^{N_{g}} \sum_{j=1}^{N_{s}} {P(i,j)j}^{2}}{N_{z}}$$

3. Gray Level Non-Uniformity (GLN)

$$GLN=\frac{\sum_{i=1}^{N_{g}} (\sum_{j=1}^{N_{s}} {P(i,j))}^{2}}{N_{z}}$$

4. Gray Level Non-Uniformity Normalized (GLNN)

$$GLNN=\frac{\sum_{i=1}^{N_{g}} (\sum_{j=1}^{N_{s}} {P(i,j))}^{2}}{N_{z}^{2}}$$

5. Size-Zone Non-Uniformity (SZN)

$$SZN=\frac{\sum_{j=1}^{N_{s}} (\sum_{i=1}^{N_{g}} {P(i,j))}^{2}}{N_{z}}$$

6. Size-Zone Non-Uniformity Normalized (SZNN)

$$SZNN=\frac{\sum_{j=1}^{N_{s}} (\sum_{i=1}^{N_{g}} {P(i,j))}^{2}}{N_{z}^{2}}$$

7. Zone Percentage (ZP)

$$ZP=\frac{N_{z}}{N_{p}}$$

8. Gray Level Variance (GLV)

$$GLV=\sum_{i=1}^{N_{g}} \sum_{j=1}^{N_{s}} p(i,j)\left( i-\sum_{i=1}^{N_{g}} \sum_{j=1}^{N_{s}} P(i,j)i \right)^{2}$$

9. Zone Variance (ZV)

$$ZV=\sum_{i=1}^{N_{g}} \sum_{j=1}^{N_{s}} p(i,j)\left( j-\sum_{i=1}^{N_{g}} \sum_{j=1}^{N_{s}} P(i,j)j \right)^{2}$$

10. Zone Entropy (ZE)

$$ZE=-\sum_{i=1}^{N_{g}} \sum_{j=1}^{N_{s}} p(i,j)\log_{2} \left( p\left( i,j \right)+\epsilon\right)$$

11. Low Gray Level Zone Emphasis (LGLZE)

$$LGLZE=\frac{\sum_{i=1}^{N_{g}} \sum_{j=1}^{N_{s}} \frac{P(i,j)}{i^{2}}}{N_{z}}$$

12. High Gray Level Zone Emphasis (HGLZE)

$$HGLZE=\frac{\sum_{i=1}^{N_{g}} \sum_{j=1}^{N_{s}} P(i,j)i^{2}}{N_{z}}$$

13. Small Area Low Gray Level Emphasis (SALGLE)

$$SALGLE=\frac{\sum_{i=1}^{N_{g}} \sum_{j=1}^{N_{s}} \frac{P(i,j)}{{i^{2}j}^{2}}}{N_{z}}$$

14. Small Area High Gray Level Emphasis (SAHGLE)

$$SAHGLE=\frac{\sum_{i=1}^{N_{g}} \sum_{j=1}^{N_{s}} \frac{P(i,j)i^{2}}{j^{2}}}{N_{z}}$$

15. Large Area Low Gray Level Emphasis (LALGLE)

$$SAHGLE=\frac{\sum_{i=1}^{N_{g}} \sum_{j=1}^{N_{s}} \frac{P(i,j)j^{2}}{k^{2}}}{N_{z}}$$

16. Large Area High Gray Level Emphasis (LAHGLE)

$$LAHGLE=\frac{\sum_{i=1}^{N_{g}} \sum_{j=1}^{N_{s}} P(i,j){i^{2}j}^{2}}{N_{z}}$$

**Gray Level Run Length Matrix (GLRLM) Features**

1. Short Run Emphasis (SRE)

$$SRE=\frac{\sum_{i=1}^{N_{g}} \sum_{j=1}^{N_{r}} \frac{P(i,j|\theta)}{j^{2}}}{N_{r}(\theta)}$$

2. Long Run Emphasis (LRE)

$$LRE=\frac{\sum_{i=1}^{N_{g}} \sum_{j=1}^{N_{r}} P(i,j|\theta)j^{2}}{N_{r}(\theta)}$$

3. Gray Level Non-Uniformity (GLN)

$$GLN=\frac{\sum_{i=1}^{N_{g}} \left( \sum_{j=1}^{N_{r}} P(i,j|\theta) \right)^{2}}{N_{r}(\theta)}$$

4. Gray Level Non-Uniformity Normalized (GLNN)

$$GLNN=\frac{\sum_{i=1}^{N_{g}} \left( \sum_{j=1}^{N_{r}} P(i,j|\theta) \right)^{2}}{{N_{r}(\theta)}^{2}}$$

5. Run Length Non-Uniformity (RLN)

$$RLN=\frac{\sum_{j=1}^{N_{r}} \left( \sum_{i=1}^{N_{g}} P(i,j|\theta) \right)^{2}}{N_{r}(\theta)}$$

6. Run Length Non-Uniformity Normalized (RLNN)

$$RLNN=\frac{\sum_{j=1}^{N_{r}} \left( \sum_{i=1}^{N_{g}} P(i,j|\theta) \right)^{2}}{{N_{r}(\theta)}^{2}}$$

7. Run Percentage (RP)

$$RP=\frac{N_{r}(\theta)}{N_{p}}$$

8. Gray Level Variance (GLV)

$$GLV=\sum_{i=1}^{N_{g}} \sum_{j=1}^{N_{r}} P(i,j|\theta)\left( i-\sum_{i=1}^{N_{g}} \sum_{j=1}^{N_{r}} P(i,j|\theta)i \right)^{2}$$

9. Run Variance (RV)

$$RV=\sum_{i=1}^{N_{g}} \sum_{j=1}^{N_{r}} P(i,j|\theta)\left( j-\sum_{i=1}^{N_{g}} \sum_{j=1}^{N_{r}} P(i,j|\theta)j \right)^{2}$$

10. Run Entropy (RE)

$$RE=-\sum_{i=1}^{N_{g}} \sum_{j=1}^{N_{r}} P(i,j|\theta)\log_{2} \left( p\left( i,j | \theta\right)+\epsilon\right)$$

11. Low Gray Level Run Emphasis (LGLRE)

$$LGLRE=\frac{\sum_{i=1}^{N_{g}} \sum_{j=1}^{N_{r}} \frac{P(i,j|\theta)}{i^{2}}}{N_{r}(\theta)}$$

12. High Gray Level Run Emphasis (HGLRE)

$$HGLRE=\frac{\sum_{i=1}^{N_{g}} \sum_{j=1}^{N_{r}} P(i,j|\theta)i^{2}}{N_{r}(\theta)}$$

13. Short Run Low Gray Level Emphasis (SRLGLE)

$$SRLGLE=\frac{\sum_{i=1}^{N_{g}} \sum_{j=1}^{N_{r}} \frac{P(i,j|\theta)}{i^{2}j^{2}}}{N_{r}(\theta)}$$

14. Short Run High Gray Level Emphasis (SRHGLE)

$$SRHGLE=\frac{\sum_{i=1}^{N_{g}} \sum_{j=1}^{N_{r}} \frac{P(i,j|\theta)i^{2}}{j^{2}}}{N_{r}(\theta)}$$

15. Long Run Low Gray Level Emphasis (LRLGLE)

$$LRLGLRE=\frac{\sum_{i=1}^{N_{g}} \sum_{j=1}^{N_{r}} \frac{P(i,j|\theta)j^{2}}{i^{2}}}{N_{r}(\theta)}$$

16. Long Run High Gray Level Emphasis (LRHGLE)

$$LRHGLE=\frac{\sum_{i=1}^{N_{g}} \sum_{j=1}^{N_{r}} P(i,j|\theta){i^{2}j}^{2}}{N_{r}(\theta)}$$

**Neighboring Gray Tone Difference Matrix (NGTDM) Features**

1. Coarseness

$$Coarseness=\frac{1}{\sum_{i=1}^{N_{g}} p_{i}s_{i}}$$

2. Contrast

$$Contrast=\left( \frac{1}{N_{g,p}(N_{g,p}-1)}\sum_{i=1}^{N_{g}} \sum_{j=1}^{N_{g}} p_{i}p_{j}\left( i-j \right)^{2} \right)\left( \frac{1}{N_{v,p}}\sum_{i=1}^{N_{g}} s_{i} \right)$$

3. Busyness

$$Busyness=\frac{\sum_{i=1}^{N_{g}} p_{i}s_{i}}{\sum_{i=1}^{N_{g}} \sum_{j=1}^{N_{g}} |ip_{i}-jp_{j}|},p_{i}\neq0,p_{j}\neq0$$

4. Complexity

$$Complexity=\frac{1}{N_{v,p}}\sum_{i=1}^{N_{g}} \sum_{j=1}^{N_{g}} |i-j|\frac{p_{i}s_{i}+p_{j}s_{j}}{p_{i}+p_{j}},p_{i}\neq0,p_{j}\neq0$$

5. Strength

$$Strength=\frac{\sum_{i=1}^{N_{g}} \sum_{j=1}^{N_{g}} \left( p_{i}+p_{j} \right)\left( i-j \right)^{2}}{\sum_{i=1}^{N_{g}} s_{i}},p_{i}\neq0,p_{j}\neq0$$

**Gray Level Dependence Matrix (GLDM) Features**

1. Small Dependence Emphasis (SDE)

$$SDE=\frac{\sum_{i=1}^{N_{g}} \sum_{j=1}^{N_{d}} \frac{P(i,j)}{i^{2}}}{N_{z}}$$

2. Large Dependence Emphasis (LDE)

$$LDE=\frac{\sum_{i=1}^{N_{g}} \sum_{j=1}^{N_{d}} P(i,j)j^{2}}{N_{z}}$$

3. Gray Level Non-Uniformity (GLN)

$$GLN=\frac{\sum_{i=1}^{N_{g}} \left( \sum_{j=1}^{N_{d}} P(i,j) \right)^{2}}{N_{z}}$$

4. Dependence Non-Uniformity (DN)

$$DN=\frac{\sum_{j=1}^{N_{d}} \left( \sum_{i=1}^{N_{g}} P(i,j) \right)^{2}}{N_{z}}$$

5. Dependence Non-Uniformity Normalized (DNN)

$$DNN=\frac{\sum_{j=1}^{N_{d}} \left( \sum_{i=1}^{N_{g}} P(i,j) \right)^{2}}{N_{z}^{2}}$$

6. Gray Level Variance (GLV)

$$GLV=\sum_{i=1}^{N_{g}} \sum_{j=1}^{N_{d}} P(i,j)\left( i-\sum_{i=1}^{N_{g}} \sum_{j=1}^{N_{d}} iP(i,j) \right)^{2}$$

7. Dependence Variance (DV)

$$DV=\sum_{i=1}^{N_{g}} \sum_{j=1}^{N_{d}} P(i,j)\left( j-\sum_{i=1}^{N_{g}} \sum_{j=1}^{N_{d}} jP(i,j) \right)^{2}$$

8. Dependence Entropy (DE)

$$DependenceEntropy=-\sum_{i=1}^{N_{g}} \sum_{j=1}^{N_{d}} P(i,j)\log_{2} (p\left( i,j \right)+\epsilon)$$

9. Low Gray Level Emphasis (LGLE)

$$LGLE=\frac{\sum_{i=1}^{N_{g}} \sum_{j=1}^{N_{d}} \frac{P(i,j)}{i^{2}}}{N_{z}}$$

10. High Gray Level Emphasis (HGLE)

$$HGLE=\frac{\sum_{i=1}^{N_{g}} \sum_{j=1}^{N_{d}} P(i,j)i^{2}}{N_{z}}$$

11. Small Dependence Low Gray Level Emphasis (SDLGLE)

$$SDLGLE=\frac{\sum_{i=1}^{N_{g}} \sum_{j=1}^{N_{d}} \frac{P(i,j)}{i^{2}j^{2}}}{N_{z}}$$

12. Small Dependence High Gray Level Emphasis (SDHGLE)

$$SDHGLE=\frac{\sum_{i=1}^{N_{g}} \sum_{j=1}^{N_{d}} \frac{P(i,j)i^{2}}{j^{2}}}{N_{z}}$$

13. Large Dependence Low Gray Level Emphasis (LDLGLE)

$$LDLGLE=\frac{\sum_{i=1}^{N_{g}} \sum_{j=1}^{N_{d}} \frac{P(i,j)j^{2}}{i^{2}}}{N_{z}}$$

14. Large Dependence High Gray Level Emphasis (LDHGLE)

$$LDHGLE=\frac{\sum_{i=1}^{N_{g}} \sum_{j=1}^{N_{d}} P(i,j)i^{2}j^{2}}{N_{z}}$$
